# Acoustic wave-induced stroboscopic optical mechanotyping of adherent cells

**DOI:** 10.1101/2023.07.29.551092

**Authors:** Thomas Combriat, Petter Angell Olsen, Silja Borring Låstad, Anders Malthe-Sørenssen, Stefan Krauss, Dag Kristian Dysthe

## Abstract

In this study, we present a novel, high content technique using an innovative cylindrical acoustic transducer, stroboscopic fast imaging and homodyne detection to recover the mechanical properties (dynamic shear modulus) of living adherent cells at low ultrasonic frequencies. By analyzing the micro-oscillations of cells we were able to simultaneously mechanotype whole populations of cells with sub-cellular resolution. The technique can be combined with standard fluorescence imaging allowing to further cross-correlate biological and mechanical information. We demonstrate the potential of the technique by mechanotyping co-cultures of different cell types with significantly different mechanical properties.

## I. INTRODUCTION

Most cells and tissues display viscoelastic properties: when subjected to an external force they will resist deformation both with (viscosity) and without (elasticity/stiffness) loss of energy. This viscoelastic behavior, quantified by the dynamic modulus, is caused by the cells’ components and relates to their type, state and disease condition which make rheological studies of biological materials attractive. The viscoelasticity of a cell varies with many cellular processes. For example, during cell division, cell stiffness changes whereby cells in the S phase of the cell cycle are generally stiffer relative to cells in the G1 phase [1]. Another set of cellular processes impacting stiffness are epithelial mesenchymal transition (EMT) and mesenchymal epithelial transition (MET) where mobile cells generally display a softer phenotype relative to stationary cells [2]. EMT and MET are seen in a plethora of natural and diseased cellular processes such as mesoderm formation during development and cancer metastasis.

The differentiation status of cells can also influence cellular stiffness as exemplified in both mesenchymal and embryonic stem cells (SC) where undifferentiated SC display lower stiffness relative to early differentiating SC [3, 4]. In addition, various pathological conditions influence the stiffness of cells. Notably, in the context of cancer, it has been shown that cancer cells are generally less stiff than normal benign cells [5–8].

The viscoelasticity of cells is determined by the main mechanical elements of the cell, the nucleus and cytoplasm. The nuclear viscoelasticity is predominantly determined by the conformation of the chromatin and the structure of the nuclear lamina [9, 10]. The main component of the cytoplasm that governs viscoelasticity is the organization of the cytoskeleton (actin filaments, microtubules and intermediate filaments) with contributions of the cortical cytoskeleton that maintains cell shape and can generate cell motion. In cells that adhere to an extracellular matrix (ECM) through focal adhesion complexes or to other cells through adherence junctions, actin stress fibers between the attachment sites are the most dynamic contributors to cytoplasm viscoelasticity [11] linked to biological functions as given by its differentiation state or disease conditions such as cancer metastasis [3, 8]. The viscoelasticity of the ECM influences the mechanical properties of adherent cells through mechanosensitive proteins in the adhesion complexes [12, 13]. In addition, temperature, pH and chemical agents and other cytoplasmic components like polymers of glycosaminoglycans and polysaccharides together with membrane tubes of the endoplasmic reticulum (ER) can also influence on cellular viscoelacticity [14–16].

Actin stress fibers remodel according to the cell adhesion to, and elastic properties of the substrate [11, 13, 16–19]. It is well established that cytoskeleton actin stress fibers have a large effect on cell’s mechanical properties [20] which can differ if the cell is adherent or suspended. For example, inactivation of contractile actin stress fibers by blebbistatin has been shown to reduce the stiffness of adherent cells by a factor of 2 [21]. Another example is an increase in stiffness with population doubling of mesenchymal stem cells that is observed in the adherent state but not in the suspended state [3]. Hence, the measurement of suspended cell stiffness does not necessarily correlate with the rheological properties of adherent cells [16]. Therefore reliable measurement of viscoelasticity of adherent cells will provide biological insights that are not necessarily possible to get on suspended cells.

At current, cell viscoelasticity of adherent cells can be determined by a spectrum of techniques including micropipette aspiration, magnetic bead rheometry, optical tweezers [22], particle-tracking microrheology [23], stretchable micropost arrays [24], stretchable PDMS substrate [25], laser Doppler vibrometer measurements of cell height during vertical vibration [26], Brillouin microscopy [27] and atomic force microscopy (AFM) [17]. AFM is the most commonly used technique combining well-controlled force, high sensitivity and spatial resolution.

All these techniques, however, are serial; they apply to one position at one cell at a time rather than being scalable to high throughput data acquisition. Since most of the techniques listed above rely on identifying measurement positions on the cell before performing the elastic or viscoelastic measurements, they are in principle also difficult to automate.

Technologies for fast measurements of the viscoelasticity of a whole adherent (2D) cell population yielding reliable measurement of the rheological properties of each cell compared to surrounding cells and to the whole 2D cell population are currently not available.

In this communication, we describe a novel, fast, precise and reliable technique to measure the viscoelastic properties of a population of adherent cells. The technique is based on a combination of three components that have not been applied to measure viscoelasticity of adherent cell before: a novel deformation transducer adapted to transmitted light microscopy, stroboscopic imaging and homodyne detection of cell strains. The combined setup, termed stroboscopic optical mechanotyping, allows unprecedented high throughput rheological measurements of adherent cells with sub-cellular resolution.

## II. RESULTS

All experiments were performed in petri dishes at room temperature on an inverted motorized microscope fitted with both a fast camera for stroboscopic imaging and a high-sensitivity camera for fluorescence (see Figure 1 and). A shear deformation stimulation was produced by custom-made focused ultrasound (US) transducers that allow traditional phase contrast and fluorescence imaging to be performed in the sonicated area. Stroboscopic imaging (see V C 2) allowed us to recover the micro-motions of the cells at the ultrasonic frequency caused by the mechanical stimulation.

### 1. Acoustic shear flow transducer

Unlike usual ultrasound transducers that need to be put at an angle with respect to the illumination to free the light path, the system used here allows for the transducer to share a common axis with the illumination. The transducer (see Figure 1 and V B 1), consisting of a piezo ring with a focusing lens surrounding a water cavity held in place by two thin glass plates, was optimized by finite difference simulations of sound propagation.

**FIG. 1.**
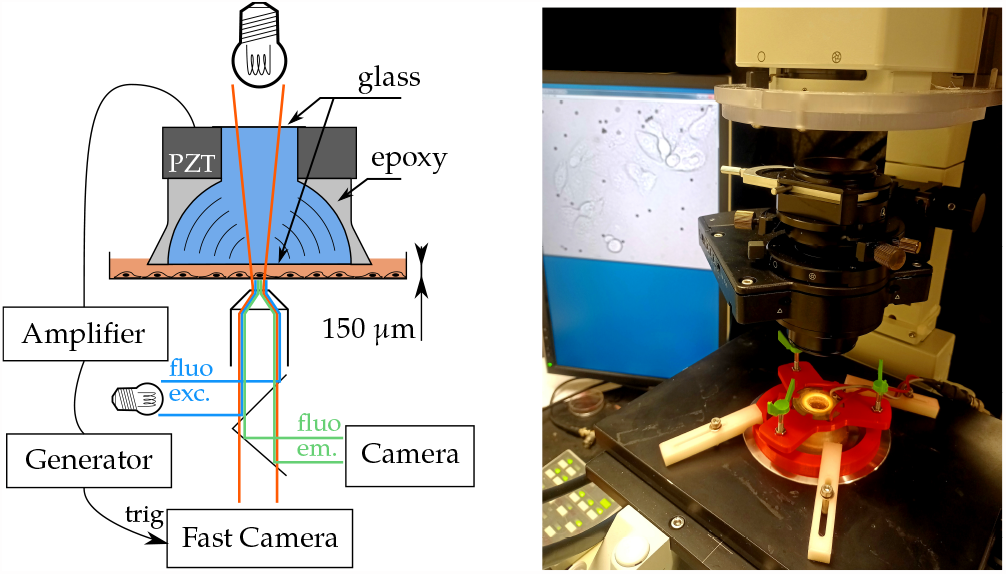
Acoustic excitation and imaging setup. Left: Acoustic tranducer consisting of piezoelectric element (PZT) with epoxy lens focusing acoustic waves through (blue) water enclosed between upper and lower glass window. The transducer is lowered into the growth media (orange) in a petri dish until the distance between the transducer glass and petri dish is about 150 μm. Cells adhering to the petri dish are deformed by shear flow in the growth media caused by the acoustic waves. The petri dish is placed on an inverted microscope with both transmitted light illumination for fast imaging of the cells while they move in the shear flow, and reflected light illumination for fluorescence imaging. A function generator generates wave pulses that are amplified and sent to the PZT and triggers the fast camera for the stroboscopic imaging. **Right:** Acoustic transducer placed in its adjustable holder (red) on the inverted microscope with the images of the cells in the petri dish displayed on the monitor behind.

To reduce acoustic impedance contrast and thus increase acoustic wave transmission, the empty space of the lens was filled with degassed water and the top and bottom parts were sealed. The transducer was subsequently suspended over a 6 cm Petri dish containing the cells to be probed (see V A) using a 3D printed holder adjustable in height and orientation.

The bottom glass plate moves vertically with the focused ultrasound wave. This motion drives an oscillating radial fluid flow with the same frequency as the acoustic wave in the fluid between the glass plate and the Petri dish where the cells adhere. This fluid flow acts on the cells with an oscillating viscous shear stress that causes a shear deformation of the cells.

### 2. Acoustic waves inducing shear deformation of cells

The signals sent to the transducer were Gaussian pulses:

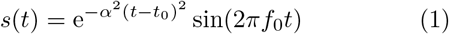

where *t* is time, *f*0 is the central frequency (*f*0 = 30 and 45 kHz for the experiments presented here), 1*/α* is proportional to the duration of the pulse (specifically, the standard deviation of the pulses is 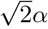) and *t*0 the time at which the maximal amplitude of the Gaussian envelope is reached. The pulses were repeated *N* times (*N* = 1000 for the experiments presented here) at a constant repetition rate *f*repet (*f*repet = 1 kHz for the experiments presented here), allowing time for the waves to fade out between the successive pulse and preventing the formation of stationary waves. The transducers were calibrated using a calibrated hydrophone placed a centimeter away from the bottom plate of the transducer (see Suppl. Fig. 8 and V B).

### 3. Stroboscopic imaging

Stroboscopic imaging (see V C 2) was used in order to recover cells’ oscillation at the US frequency while imaging at a lower frame-rate, thus improving the field of view that could be probed in a single experiment. The images, taken through a 20X objective, are focused 2-4 micrometers above the substrate where the cells adhere. The images record the positions in the image plane (xy-plane) of the cells and their internal structure resolved by phase contrast. The displacement of the cell structures in the image plane allows calculation of the in-plane deformation of the cells

#### A. Image analysis

An overview of the data analysis process is shown in Fig. 3B and the different steps are detailed below.

##### 1. Segmentation

Segmentation was performed on the images acquired with the fluorescence imaging system (see V C 1). All fluorescent images were first averaged to increase the signal-over-noise ratio. Long-range heterogeneity (background) in the illumination was computed using a Gaussian blur with a kernel size that is half the images’ size. This background was subtracted from the averaged image and the result was segmented to identify biological material in the field of view by standard threshold.

In experiments with several fluorescence channels, i.e. with several cell types mixed, the intersecting areas between the different masks were removed from the analysis. This corresponded to areas where the two cell lines were in contact and concerned less than 5 % of the data.

##### 2. Homodyne stress estimation

Particle Tracking Velocimetry (PTV) was performed on the tracer particles (see V B 3) located in the same focal plane as the biological cells studied. These beads rest on the bottom of the dish.

A homodyne detection was performed on these displacements by performing a fast Fourier transform (FFT) on the averaged particle displacements over time and keeping only the complex coefficient at the central frequency:

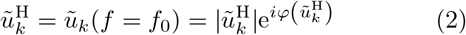

where *ũk* is the temporal Fourier transform of the averaged beads displacements along the *k x, y* axis, *φ* (*ũk*) is its phase and *i* is the imaginary unit. Note that in all of this paper the H superscript denotes an homodyne detection, *i*.*e*. that only the Fourier components at the carrier frequency *f*0 where kept.

This homodyne detection, in a similar fashion to lockin amplifiers, allows to improve the signal-to-noise ratio between one and two orders of magnitudes by filtering out all the noise present at frequencies other than the carrier frequency *f*0.

By considering the fact that the beads used for this estimation were lying on the bottom, the homodyne complex stresses applied to the cells in both directions were approximated as:

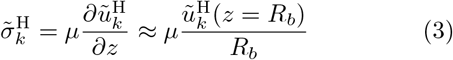

where *μ* is the dynamic viscosity of media, approximated as the one of water at room temperature and *Rb* the beads radius (see V B 3). More details are given in V D 1.

##### 3. Homodyne cell motion estimation

Digital image correlation (DIC) was used on every image to obtain displacements δ*x*(*t*) and δ*y*(*t*) in the *x* and *y* directions with respect to a reference image (see Fig. 2 top).

**FIG. 2.**
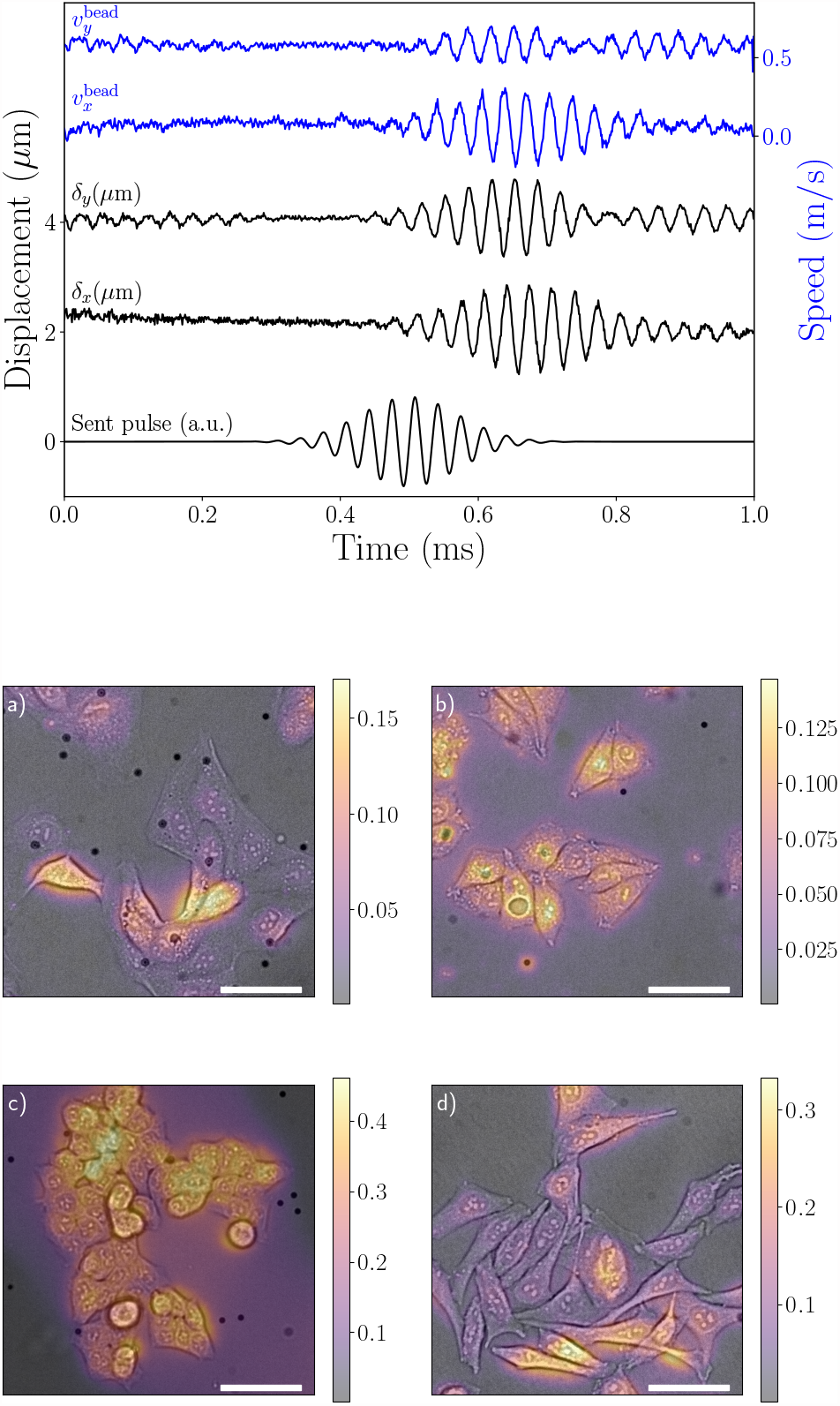
Top: Typical cell displacements of cells and beads. Cell displacement δ*k* (black) recovered by DIC (before homodyne filtering) and bead speeds *vk* (blue) recovered by PTV in the *k x, y* directions versus time. The displacements are averaged over the whole image. The signal sent to the generator is depicted in arbitrary units at the bottom for comparison. The curves are vertically shifted for visibility. **Bottom: Examples of homodyne displacement amplitudes**, 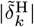, (color coded) of different adherent cell types overlayed phase contrast images of the cells. a) MDCK, b) MelJuSo, c) HTC116, d) HeLa. Beware that the color bars are scaled individually. The scale bar is 50 μm.

Analogous to the flow field estimation (see II A 2) homodyne detection was performed on the cell motion maps. This yields:

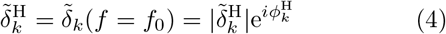

where 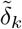 is the temporal Fourier transform of the displacements in the *k* ∈ {*x, y*} directions.

This leads to pixel-wise homodyne amplitude 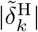 and phase 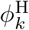 maps (see Figs. 2 and 3).

**FIG. 3.**
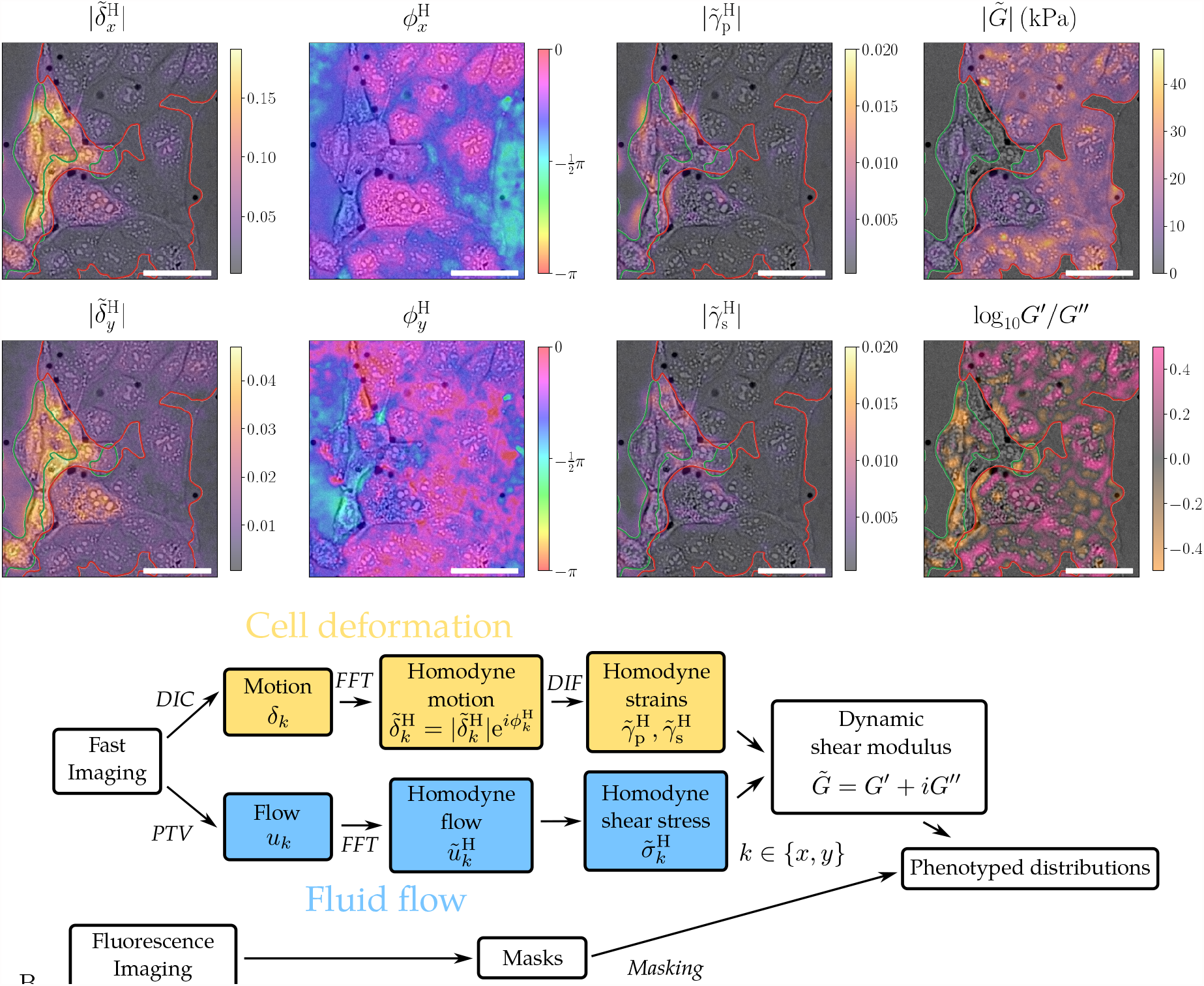
A: Results from each step in the analysis workflow. The images are phase contrast images for a co-culture of MelJuSo (green contours) and MDCK cells (red contours) with the values of different measures overlayed in colour. **Column 1 and 2:** The digital image correlation (DIC) and FFT yield homodyne displacement amplitude 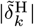 and phase 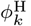. One observes that MelJuSo cells move more and have a larger phase shift than the MDCK cells. **Column 3:** Differentiation of the homodyne displacements yield the homodyne pure 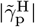 and simple 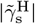 strain amplitudes (see eqs. (5) and (6)). One observes that MelJuSo cells are more strained than the MDCK cells. **Column 4:** The fluid stress on the cells divided by these strains yield the dynamic shear modulus 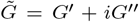. Top: The shear modulus amplitude 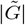 is the common measure of cell viscoelasticity. Bottom: The ratio between the storage modulus *G′* and loss modulus *G″* is the ratio between elastic and viscous components. One observes that MelJuSo cells are softer and more viscous whereas MDCK cells are stiffer and more elastic. Beware that the color bars are scaled individually. The black, circular dots are the tracer beads used to measure the fluid flow. Scale bar is 50 μm. The shown images cover 1*/*20 of the full field of view. **B: Workflow diagram of the analysis process**. Analysis of the fast imaging: The fluid flow is measured by PTV and the shear stress is calculated from the vertical flow gradient. The cell deformation is quantified by digital image correlation (DIC) and the gradients (DIF) in displacement yield the strain. In both paths the accuracy of the measurement is greatly increased by homodyne detection, that is, using FFT to only sum over changes at the imposed acoustic frequency. The shear modulus is the ratio of the resulting homodyne stress and strain. Fluorescence images were used to produce masks to identify position of cells and to differentiate between cell types. Comparison of shear moduli of different cell types in a mixed cell culture allowed identification of mechanotypes.

##### 4. Homodyne strains computation

Homodyne pure 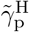 and simple 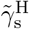 shear strains are computed from the homodyne displacements maps 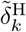 (see V D 2):

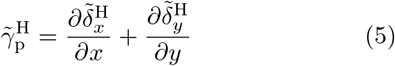

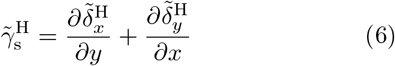

These homodyne shear strains are complex numbers that carry information about both the amplitude of deformation around the rest state (through the absolute value) and the phase difference of these deformations with respect to the external forcing.

##### 5. Shear modulus

The dynamic shear modulus of a material is defined as the ratio between the shear stress and the shear strain:

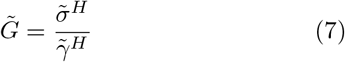

The exact manner how this is calculated by summation over the different spatial components is given in the Materials and Methods section. The dynamic shear modulus 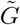 is a complex number that can further be decomposed in terms of the storage modulus *G′* representing the elastic component and the loss modulus *G″* representing the viscous component:

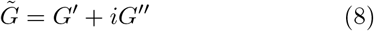

The novel instrument and method of analysis have been used on several different cell types that all show shear displacements that are characterized by a good signal-to-noise ratio. In Fig. 2 we show homodyne displacement amplitudes of 4 different cell lines, Madin Darby canine kidney cell line MDCK, human melanoma cell line MelJuSo, human colorectal carcinoma cell line HTC116 and cervical cancer cell line HeLa (see section V A). The shown displacement maps are the results of the homodyne detection on digital image correlation performed on the movies available in the Supplementary information. In the following, we will focus on the detailed analysis results of two cell types, MelJuSo and MDCK, to display the ability of the method to distinguish the cells by mechanotyping.

#### B. Motion, strain and mechanotyping at the cellular level

With a single ultrasound burst our method yields bidimensional motion, strains and dynamic modulus maps of living cells in fields of view up to the size of a colony, up to 733 × 676 μm^2^ for the experiments presented here. Fig. 3, shows the intermediate results of the image analysis up to the final spatial distribution of elastic and viscous properties in part of a co-culture of two cell types. The homodyne displacement amplitudes 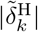, are much larger for the cells on the left in the image than those on the right. The pink phase amplitudes 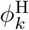, displayed by the cells with small displacement amplitudes indicate a small phase lag with respect to the fluid motion and the purple and blue phase amplitudes indicate that the regions with higher displacement also have a larger phase lag. The pure and simple shear strain amplitudes 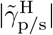, calculated according to Eqs. (5) and (6) are largest for the cells with large displacement amplitudes. The fluid flow field calculated from the motion of the beads is homogeneous over the field of view in Fig. 3. The flow field is used to calculate the fluid shear stress and the ratio of shear stress and shear strain yields the dynamic modulus 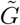 (see Eq. (20)). The absolute value of the dynamic modulus 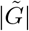 and the ratio of the real *G ′* and imaginary *G ″* parts are shown to the right in Fig. 3. As expected, the least moving cells are stiffer (higher dynamic modulus) and the cells moving the most are softer. One may also observe some internal variation in the viscoelasticity of the cells and the intracellular variations are especially visible for the ratio between the storage (elastic) and loss (viscous) modulus.

When studying the rheology of complex materials it is customary to display the ratio of storage over loss modulus as a function of frequency. In Fig. 3 right we display the map of the storage to loss modulus ratio for the same field of view as before. The map has a consistently high signal-to-noise ratio and displays that extending the method to performing frequency sweeps can yield significant information on the viscous versus elastic behavior of the cell cytoskeleton.

#### C. Different cell types display different mechanotypes

For both frequencies studied, 30 and 45 kHz, mechanotyping using our stroboscopic optical apparatus was performed on mixed and separated populations of fluorescent labelled MelJuSo and MDCK cells. The experiment on mixed populations was repeated three times (technical replicates on different dishes) for each frequency. Additionally, both cell types were tested once in separated cultures for both frequencies (30 kHz and 45 kHz).

In Fig. 4 we show the dynamic modulus of a co-culture of MDCK and MelJuSo cells. The fluorescence image of the same field of view was used to segment the image into three categories: petri dish (dark background), MelJuSo cells (green contours) and MDCK cells (red contours). The right side of Fig. 4 shows the statistics of dynamic modulus 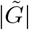, storage modulus *G ′* and loss modulus *G ″* for the imaged cells at 30 kHz. The box plots are calculated using the DIC-box (see V D 2 and following) as the statistical unit with global cell masking based on the fluorescence signal. One observes that even though many MelJuSo cells are stiffer than certain MDCK cells, the population statistics using the MannWhitney U-test clearly separates the two cell populations. The dynamic moduli with standard error of the mean (SEM) are 4.40 *±* 0.03 kPa for MelJuSo and 6.29 *±* 0.05 kPa for MDCK for this experiment.

**FIG. 4.**
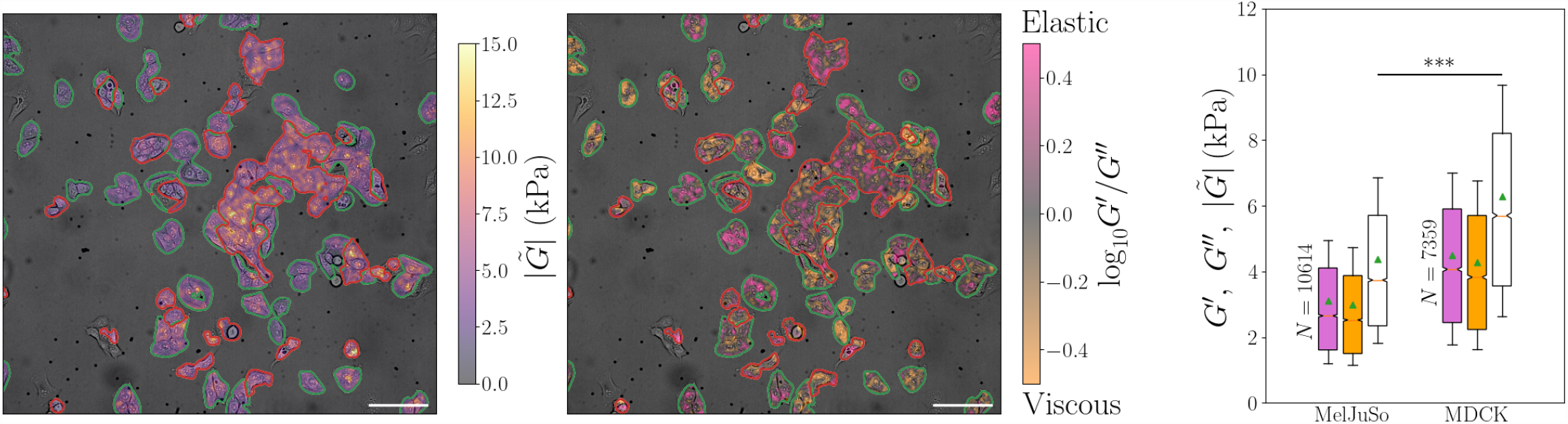
Cell dynamic shear modulus, 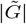 and the elastic, *G′*, and viscous, *G″* components for a mixed cell culture of MelJuSo and MDCK . MelJuSo and MDCK cells are identified by correlative fluorescence microscopy and contoured in green and red, respectively. The scale bar in both images is 100 μm. **Left:** Dynamic modulus 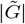 map of cells overlayed phase contrast image. One readily observes that most MDCK cells are stiffer than most MelJuSo cells. **Middle:** Map of the ratio of storage and loss moduli log10 *G′/G″* overlayed phase contrast image. The ratio *G′/G″*, that is whether the cells are more elastic or viscous on the other hand, does not depend on the cell type. **Right:** Boxplot showing statistics of dynamic modulus 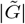 (white), storage modulus *G′* (purple) and loss modulus *G″* (orange) from the field of view shown in left and middle. The orange line corresponds to the median of the distributions and the green triangle is the mean. The statistical unit used is the DIC-box and statistical significance is obtained by a Mann-Whitney test with *** corresponding to a p-value *<* 0.001. Measurements were performed at 30 kHz

In Fig. 5 we summarize all the experiments on MelJuSo and MDCK, using DIC-box based statistics. One may observe that MDCK have higher moduli than MelJuSo and that the storage (elastic, in purple) modulus is larger than the loss (viscous, in orange) modulus and the absolute value of the dynamic modulus (white) is always largest. It can also be seen that the mechanical properties of MDCK are similar for the two probed frequencies whereas for MelJuSo, all components of the dynamic modulus show an increase at 45 kHz compared to 30 kHz, which could indicate that the main component responsible for the mechanical behavior of cells is different for these two cell lines. The mechano-contrast between these two cell lines is greater at 30 kHz than at 45 kHz, hence separating these two cell lines using mechanical properties alone is better performed at 30 kHz. Frequency scanning could help finding the optimal probing frequency for separation but is beyond the main scope of this communication.

**FIG. 5.**
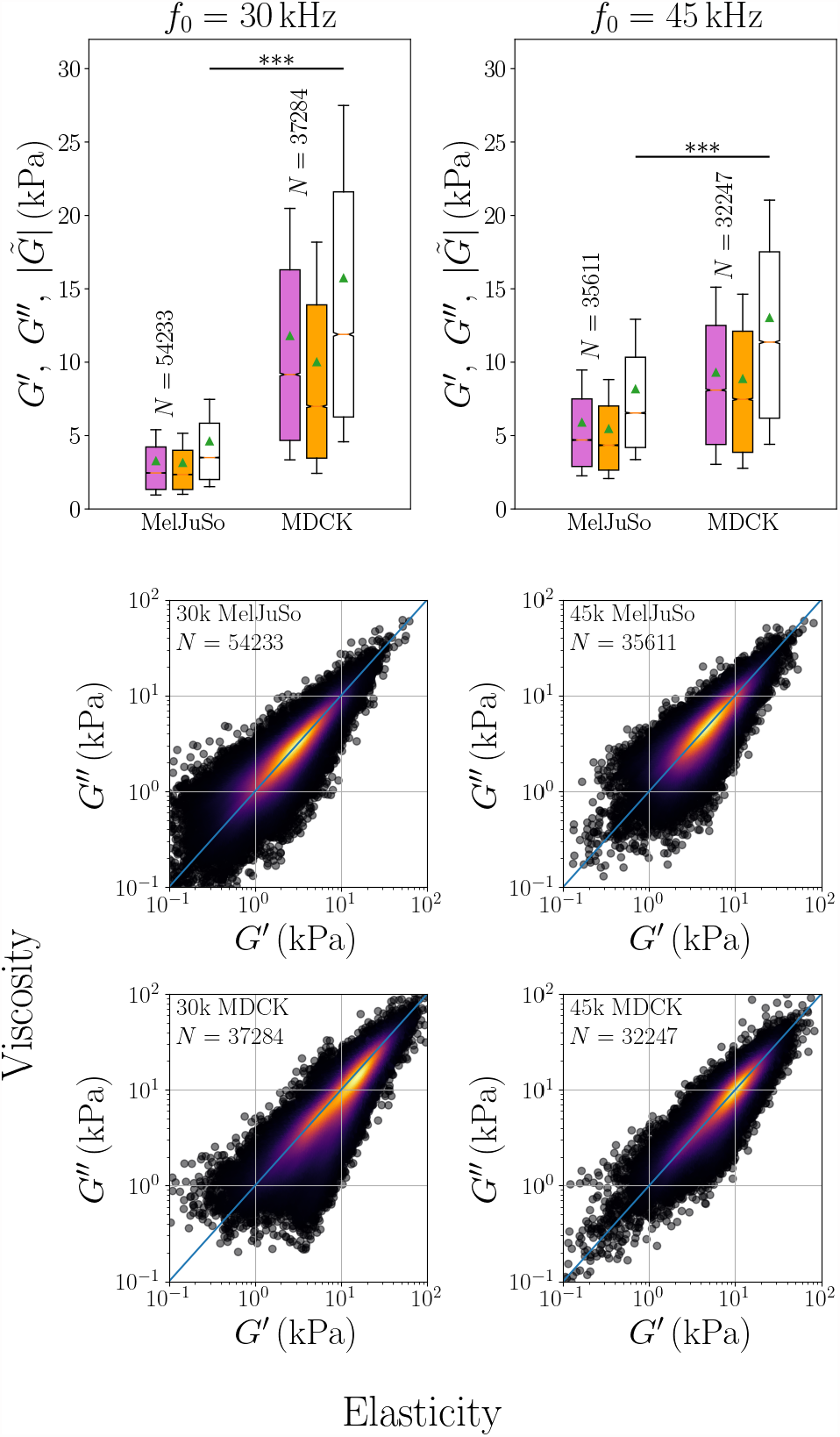
Top: Dynamic shear moduli for the two cell types cultured both together (like in Figure 4) and separately. Cell viscoelasticity (dynamic shear modulus) 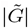 (white), elastic component (storage modulus) *G′* (purple) and viscous component (loss modulus) *G″* (orange). Statistics are performed using the DIC-box as the statistical unit. Outliers are not shown. Each shown distributions are composed of three repetitions on separate dishes with mixed cell types and one with one cell type alone. **Bottom: Elastic**, *G′*, **and viscous**, *G″*, **components are strongly correlated for all cell types probed in this study:** scatter distributions of *G″* as a function of *G′* for populations of two different cell types (MelJuSo and MDCK) at two different frequencies (30 kHz and 45 kHz). Each scatter represents one DIC-box. Color depicts the local density of scatters, computed as a Gaussian-kernel density estimate and the line is *G″* = *G′*.

#### D. DIC-based vs. cell-based statistics

When analysing statistical variation in elastic properties of cells it is natural to find the average for each cell and then calculate and display the inter-cell distribution. This statistical analysis, that we will label cell based statistics (CS), requires segmentation of each individual cell in the image, a task that is usually difficult to perform on colony-forming cells. Our approach has instead been to measure the elastic properties for every group of pixels, the DIC-box (2.57 *×* 2.57 μm^2^, see V D 2), used to perform the digital image correlation. The values measured for each DIC-box are then used to calculate the statistical distribution that we will label the DIC-box based statistics (DS). In the case of the mixtures of MelJuSo and MDCK we only need to use global cell masking instead of individual cell identification to perform the DIC-box based statistics (DS).

To assess the pertinence of this approach, cell based segmentation was manually performed on one experiment on a mixed population of MDCK and MelJuSo and its statistics were compared to the ones recovered using DIC-boxes as statistical units. Suppl. Fig. 7 shows the resulting boxplots using cells (CS) or DIC-boxes (DS) as statistical units. It can be seen that means, medians, quartile positions and standard deviations are very similar with both techniques showing that DIC-box based statistics capture the same information as cell based statistics while significantly simplifying the analysis method. The main difference between the cell based (CS) and DIC-box based (DS) statistics is that the cell based standard error of the mean (SEM), which is indicated by the height of the notches on the boxes is only biologically meaningful for the cell based statistics.

The agreement between the boxplots using cell-based and DIC-based statistics (Suppl. Fig. 7) is the basis for using this instrument for high content screening of viscoelasticity of cell colonies. The DIC-based calculation can be fully automated since it avoids error-prone segmentation that often requires human intervention during analysis.

#### E. Intracellular variations, storage and loss modulus

The attainable spatial resolution of the displacements, strains and moduli is principally dictated by the global optical resolution of the system and the size of the comparison box used for the DIC (DIC-box). The spatial resolution is 2.57 μm for the experiments presented here. This resolution is smaller than the typical size of adherent cells used in *in vitro* experiments, giving access to both the imposed strains and mechanical properties at the sub-cellular level. It is worth noting, as it can be seen in Fig. 3 (left), that cells’ nuclei are generally associated with strong homodyne displacement components. The viscoelasticity, however is related to the spatial derivatives of the displacement. When we average over the dynamic modulus of many cells from the edge and towards the center (see Fig. 6) we find that the cells are softer in the center (where the nuclei tend to be) than at the edge of the cells. This conforms well with AFM measurements of cell rheological properties. Abidine et al. [28] have compared the storage and loss moduli on top of the nucleus, at the site of the nucleus and at the cell edge and found that at low frequency the modulus decreases markedly from the edge to the nucleus. At higher frequencies (up to 300 kHz) the difference persists but becomes smaller.

**FIG. 6.**
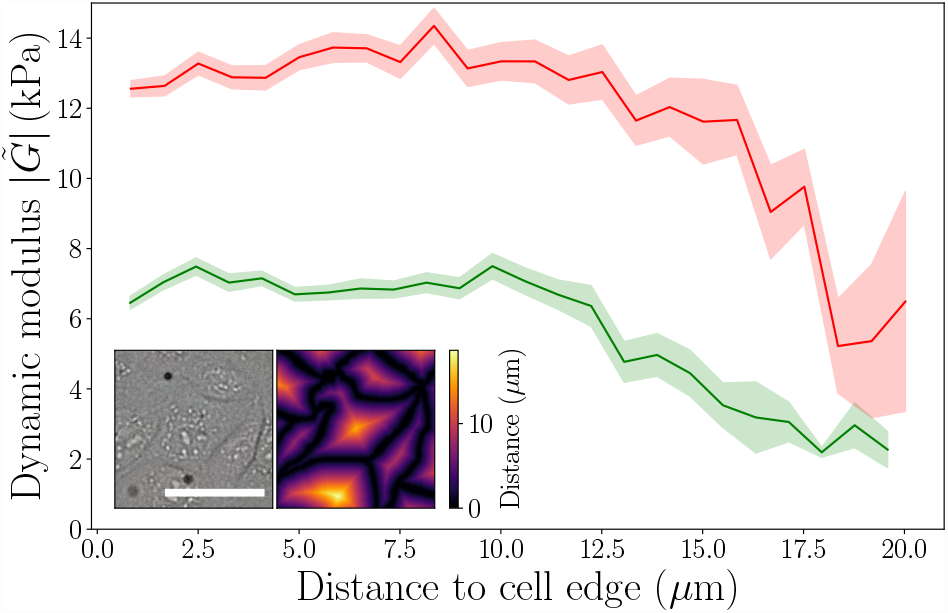
Viscoelasticity as function of distance from the cell’s edge for MelJuSo (green 68 cells) and MDCK (red 80 cells) cell types. Both cell types are softer in the centre (in the vicinity of the nucleus). Performed on a single experiment with a mixed population of MelJuSo and MDCK with individually labelled cells. Inset: example of phase contrast image and distance to cell edge metric computed on the same FOV. The scale bar is 50 μm.

**FIG. 7.**
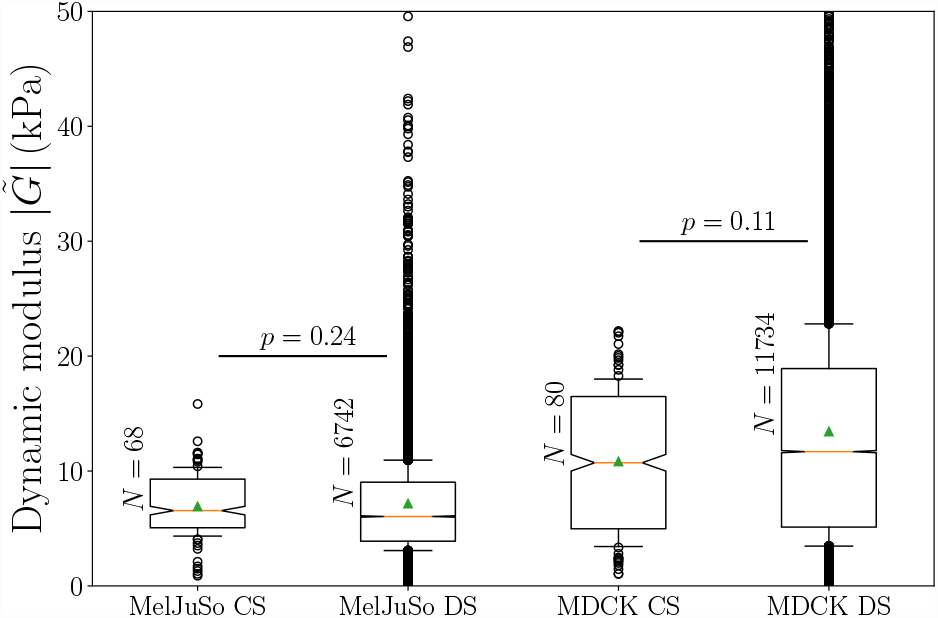
Dynamic shear modulus 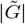 boxplots calculated using cells (CS) and DIC-box (DS) as the statistical unit. Data are extracted from one experiment done on a mixed population of cells (MelJuSo and MDCK). The number of statistical units, *N*, for each boxplot is a factor 100 larger for DIC-box than for cells. Despite this the two distributions for each cell type have very similar median and standard errors showing that DIC-boxes can be used as the statistical unit. The displayed *p*-values are the results of a Mann-Whitney U test.

**FIG. 8.**
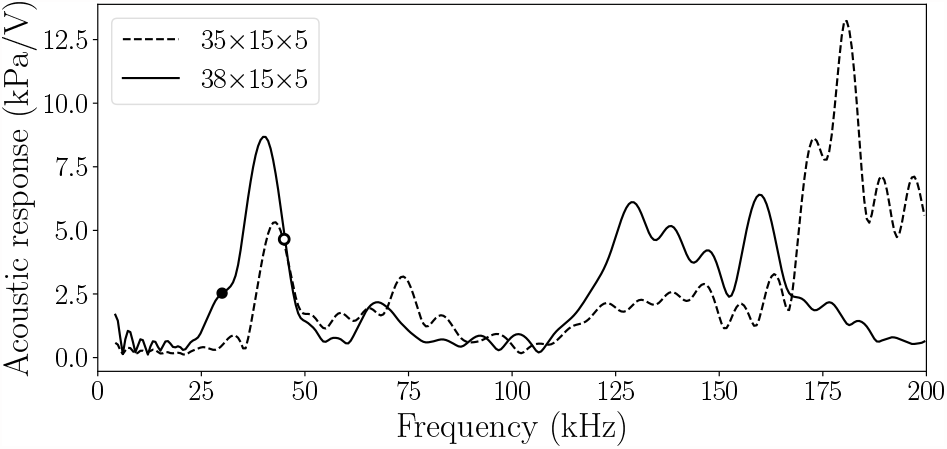
Calibration of the two transducers used in this study. The dots denote the frequencies used for each transducer.

The large number of DIC-box based measurements allow us to study new relations. Fig. 5 (bottom) shows the distributions of the storage, *G′*, versus loss, *G″*, moduli for the two cell types at the two frequencies used. The large spread in values of the moduli represent the large differences between different cells and the large differences between different parts of cells. It is very striking, however that the storage and loss moduli are proportional and that most frequently the two are more or less equal. This is a type of rheological study we have not encountered before and will only attempt a hypothesis opened to further study. The equality of *G′* and *G″* hints that the structure of the viscoelastic cytoskeleton network in the cell is roughly the same for both weak and strong cells/parts of cells.

#### F. Comparison to other techniques

Although many techniques have been developed for measuring mechanical properties of adherent cell, AFM remains the gold standard. AFM measurements of microrheology can be done at frequencies up to 100 *–* 300 Hz with a standard AFM [29–31] and up to 100 kHz with a specialized highfrequency AFM [32]. Rolling magnetic bead microrheology has been used up to 1 kHz [33]. AFM measurements typically report the standard error of the mean (SEM) and the sample standard error is of the order 100-200%, which is similar to the sample standard error in our measurements. Since our technique easily measures the moduli of 70-80 cells simultaneously the SEM (using cell based statistics) is as low as 10-20% (see Suppl. Fig. 7).

The storage modulus of cells reported in measurements by AFM follow roughly a power law behavior with frequency with exponent 0.2-0.4, for epithelial cells typically starting at a few hundred Pa at low frequency to around 1 kPa at 100 Hz and around 10 kPa at 100 kHz. Typically the loss modulus is lower than the storage modulus at low frequencies and they approach each other at high frequencies [29–32, 34]. Hence, the moduli measurements in this study agrees very well with the AFM measurements on similar cell types.

## III. OUTLOOK

We have presented a new, high content mechanotyping technique. Using a novel cylindrical acoustic transducer, stroboscopic fast imaging and homodyne detection the technology recovers elastic and viscous moduli of live adherent cell cultures. The entire field of view is probed simultaneously at sub-cellular resolution yielding high statistical significance of the results from a single experiment burst of 1 s duration.

The optical resolution in this proof-of-concept, was 2.57 μm, allowing sub-cellular components to be identified, whereby the viscoelasticity of the nucleus can be differentiated from that of the cytoskeleton. The resolution and the probed area are only limited by the imaging system, hence, higher resolution and field of view (FOV) should be attainable by the use of higher or lower magnification lenses given that enough light is provided to attain the short exposure times required by the stroboscopic measurements. In that respect, the use of cylindrical hollow acoustic lenses allows good conditions for transmission illumination imaging of the area being insonicated. Finally, this technique is evolving in the low-ultrasonic range and provides viscoelastic measurements at frequencies not attainable by standard AFM elastography. The lower boundary of this range is limited by the possibility to build a practical hollow transducer for audible frequencies, but at lower frequencies microfluidic approaches to shear flow can be used. The high frequency boundary of the technique is limited by image acquisition (exposure time and illumination). We foresee that the technology can be used to explore a variety of biological processess, including epithelial mesenchymal transition (EMT), extravasation, epithelial barrier functions and others, in a complex 2D cellular landscape with sub-cellular resolution.

## IV. ACKNOWLEDGEMENTS

The work has received funding from the University of Oslo Life Science Program “Convergence Environment” and the Research Council of Norway “Centre of Excellence” Scheme (project number 262613). The authors gratefully acknowledge the long term loan of a high speed camera from the CoE PoreLab.

## NOMENCLATURE

*f*_0_: Carrier frequency of the ultrasonic signals
*u*_*k*_: Flow displacement fields alongside *x* and *y* directions
*ũ*_*k*_: Fourier transforms of the *u_k_*
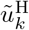: Homodyne flow displacement fields: 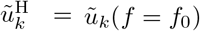
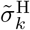: Homodyne stresses alongside *x* and *y* directions
μ: Dynamic viscosity
δ_*k*_: Cell motion fields alongside *x* and *y* directions
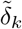: Fourier transform of δ_*k*_
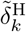: Homodyne cell motion fields: 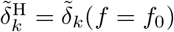
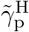: Homodyne pure strain field
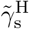: Homodyne simple strain field
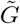: Dynamic/viscoelastic shear modulus
*G′*: Elastic modulus
*G″*: Viscous modulus

## V. MATERIAL AND METHODS

### A. Cell culture

HCT 116 (human colon cancer) cells (American Type Culture Collection (ATCC - CCL-247) were cultured in McCoy’s 5A medium (Sigma-Aldrich), HeLa (human cervical carcinoma) (ATCC CCL-2), MelJuSo (human melanoma) [35] and the MDCK (Madin-Darby canine kidney strain II) cells (ECACC - 00062107) were cultured in complete Dulbecco’s Modified Eagle Medium (DMEM, Sigma-Aldrich). All cells were grown in the precence of 10 % FBS (Sigma-Aldrich) and 1 % penicillin/streptomycin (Sigma-Aldrich) at 37 °C in a humidified atmosphere containing 5 % CO2. For ultrasound (US) experiment cells were grown in 6 cm dishes (Nunc, cat. no: 150288) and during experiments (outside of the cell incubator), cells were kept in “CO2 Independent Medium” (Thermo Fisher Scientific) supplemented with 10 % FBS and 1 % penicillin/streptomycin.

To facilitate segmentation during image analysis, cells were labeled with a fluorescent dye prior to US treatment. Labeling was performed by incubating the cells for 30 min in serum-free DMEM containing either 1 μM CellTracker Green CMFDA or 10 μM CellTracker Orange CMTMR (both from Thermo Fisher Scientific). To prepare a mixed culture of the two cell lines, MelJuSo and MDCK cells were separately labeled with CellTracker Green CMFDA or CellTracker Orange CMTMR, respectively. Next, the labeled cells were trypsinized, mixed in a 1:1 ratio, plated out in 6 cm dishes with DMEM and cultured for 24 hours before US treatment.

### B. Mechanical stimulation

#### 1. Annular transducer

The two transducers used in this study are based on piezo rings (PZT 4, outer diameter *×* inner diameter × thickness = 35*/*38 *×* 15 *×* 5 mm3 - PiezoElements) fitted with a semi-spherical acoustic lens. After soldering the electrodes on the piezo ring, the lens was cast on the ceramic from a mold accommodating the ring with degassed epoxy resin. The lens was made of clear epoxy resin (Araldite 2020) and the mold made of cured silicon using a replication kit (Smooth-On) from an initial 3D print (Ultimaker 2+, 100 μm vertical layer size).

To reduce acoustic impedance contrast and thus increase acoustic waves transmission, the empty space of the lens was filled with degassed type-II water and the top and bottom parts were sealed using UV glue (Express Glas Casco) with 150 μm thick glass coverslips of diameter 20 mm and 50 mm respectively. Special care was given to the absence of air bubbles in the water filling the empty space of the lens to ensure consistent acoustic properties and a light path for transmission imaging (see V C 2) comporting only elements orthogonal to the direction of the illumination.

The transducer used was suspended over the 6 cm Petri dish containing the cells to be probed (see V A) using a 3D printed holder adjustable in height and orientation. The bottom of the transducer was set parallel to the bottom of the dish at a known distance of 150 μm. This was done by interferometry between the bottom of the transducer and a 50 mm diameter, 150 μm thick cover slip placed in an empty dish for the initial alignment. This alignment procedure was performed only once for a series of experiments, considering that the geometrical properties of the different dishes used were identical.

#### 2. Signal generation and amplification

The signals sent to the transducer were generated by the arbitrary wave generator of a TiePie Handyscope HS5 (TiePie Engineering) and then amplified by an AE Techron 7228 broadband amplifier set at maximal, calibrated, gain. The signals used were Gaussian pulses of the equation:

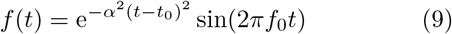

where *f*0 is the central frequency (*f*0 = 30 and 45 kHz for the experiments presented here) and *t*0 the time at which the maximal amplitude of the Gaussian envelope is reached. In the frequency domain, these pulses are gaussians centered around the central frequency *f*0 and of standard deviation 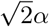 (*α* = 10 kHz for the experiments presented here). The pre-amplified amplitude of these pulses was varied from 5 to 20 Vpp leading to a driving voltage of up to 400 Vpp of the piezo. The sampling frequency of the signal was set to fifty times the central frequency *f*0 of the generated pulse.

The pulses were repeated *N* times (*N* = 1000 for the experiments presented here) at a constant repetition rate *f*repet (*f*repet = 1 kHz for the experiments presented here), allowing time for the waves to fade out between the successive pulse preventing the formation of stationary waves. Each pulse was centered in the repetition period, hence: 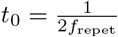.

The transducers were calibrated using a calibrated hydrophone (Type 8103 - Brüel & Kjær) placed a centimeter away from the bottom plate of the transducer. The hydrophone and the bottom plate of the transducer were immersed in a tank filled with tap water. Gaussian pulses (see Eq. 9) were generated by the arbitrary wave generator of the HS5 and sent to the transducer after being amplified. The frequency range attainable by the hydrophone (1 *–* 200 kHz) was scanned using 150 different central frequencies *f*0. For each frequency, the pulse was repeated 50 times and the signal from the hydrophone was recorded using the first acquisition channel of the TiePie. Each pulse was kept short enough so that the signal of interest did not overlap with the echoes caused by the finite dimensions of the tank. The Fourier transforms of the acquired signals were renormalized by the Fourier transforms of the original pulses and using the calibration data from the hydrophone manufacturer (see Suppl. Fig. 8).

#### 3. Tracers

Before the experiment, the cell medium was seeded with non-fluorescent tracers (Polyscience Inc. - Polybead) of a radius *Rb* = 1.5 μm. The maximum velocity attained by these beads in experiments was measured to be of the order of 1 m/s, thus the Stokes number of these particles was in the order of 10*–*2 which ensures that they are following the flow accurately.

### C. Imaging

All experiments were performed at room temperature on an inverted motorized microscope (IX81 - Olympus) controlled with μManager [36]. The microscope was fitted with both a fast camera for stroboscopic imaging and a high-sensitivity camera for fluorescence (details below). These two cameras were mounted on the side and the eye-port, respectively. Fluorescence and stroboscopic imaging were performed sequentially. Stroboscopic imaging was performed right after the fluorescence acquisition on the same field of view.

#### 1. Fluorescence imaging

The fluorescence imaging was performed with a lownoise camera (Zyla sCMOS Andor) controlled alongside the microscope with μManager [36]. Illumination was provided by a broad-spectrum mercury lamp (U-LH100HG - Olympus) before being fed to standard fluorescence cubes (U-MNIBA3 and U-MWG2 - Olympus). The intensity of the light was adjusted by using neutral density filters with optical densities of 0.2 or 0.6.

For the experiments presented here, fluorescence images were acquired prior to US stimulation at 12 bits depth. Usually, ten images for each channel were acquired and later averaged to increase the signal-to-noise ratio.

#### 2. Stroboscopic imaging

Stroboscopy is a technique allowing to recover fast periodic motions by imaging them at a lower frequency, close to or at an integer divider of the phenomenon frequency. When this is done with a short enough exposure time, the original motion is aliased at lower frequencies, appearing slower or even stationary, as it can be seen for example in movies where the wheels of vehicles appear to move slower or in the opposite direction than they actually do.

According to the Nyquist-Shannon theorem, imaging a cell’s motion at ultrasonic frequencies requires an imaging speed of more than two times the ultrasonic frequency. For the experiments presented here, where the highest ultrasonic frequency used was 45 kHz, imaging at, at least 90 kfps would have to be needed to recover the motion occurring at the US frequency. On modern fast cameras, such framerates can usually be accessed by reducing the captured field of view or binning, leading to a smaller probed material and/or lower resolution. Using stroboscopy, a large field of view can be recorded using a lower framerate while still recovering the fast motion occurring at US frequency.

Here, stroboscopic imaging was performed using a fast camera (Fastcam mini WX - Photron) under phase contrast illumination provided by the microscope builtin lamp at full power through the transducer (see V B 1). The frame acquisition was triggered by an external generator (Tektronix - AFG1062) which was clock-synchronized with the arbitrary waveform generator of the TiePie (see V B 2) that was used for signal generation. Frame acquisition rate *f*strob was set so that the equivalent of a full pulse is imaged during the repetitions, hence:

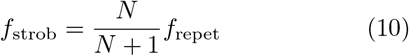

where *N* is the total number of pulses and *f*repet the repetition rate of the pulses (see V B 2). Following this, one image is acquired for each pulse leading to a minimal sequence of *N* images for complete motion recovery. In practice, a greater number (typically *N*aq = 3*N/*2) of images are acquired in order to be used as reference images. Both the camera and the triggering generator are triggered upon start by the TiePie, ensuring no jittering between the experiments.

The exposure time of the camera was set to the lowest value yielding usable images and was always under 1*/*200 ms. As a consequence, the exposure time was set to be at least four times smaller than the period of the stimulation (1*/f*0) ensuring good stroboscopic conditions. Post-experiment reconstruction was done by considering that adjacent frames are to be separated by a constant amount of time δ*t* = 1*/f*strob *–* 1*/f*repet.

### D. Data analysis

#### 1. Homodyne stress estimation

Particle Tracking Velocimetry (PTV) was performed using FAST (https://github.com/tomcombriat/FAST) on the tracer particles located in the same focal plane as the biological cells studied. These beads rest on the bottom of the dish. As the flow surrounding the cells was not homogeneous, a spatial flow estimation was performed. This was done by dividing the FOV into *n × n* smaller regions. The choice of *n* was done so that every sub-FOV would contain at least five particles during the whole duration of the experiment. The displacements of the particles in each of these areas were averaged over the particles for both directions *x* and *y* leading to displacement fields *uk, k* ∈ { *x, y*}. A homodyne detection was performed on these displacements by performing an FFT on the averaged particles displacements over time and keeping only the complex coefficient at the central frequency:

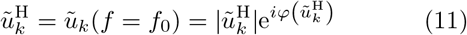

where *ũk* denotes the Fourier transforms of *uk*. Each 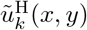 was associated with the center of one of the aforementioned areas. An estimation of the oscillating flow field was finally obtained by linear interpolation between these centers. This leads to the amplitude 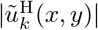 and phase 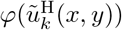 of the oscillating flow field cells were subjected to at the carrier frequency *f*0.

By considering the fact that the beads used for this estimation were lying on the bottom (ie. their center is at *Rb* with respect to the bottom of the dish) the homodyne complex stresses applied to the cells in both directions were approximated as:

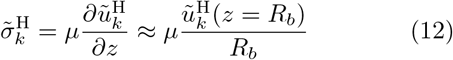

where μ is the dynamic viscosity of media, approximated as the one of water at room temperature.

#### 2. Homodyne cell motion estimation

Cell motion maps δ*k*(*x, y*) were extracted, pixel-wise for the horizontal (*x*) and vertical (*y*) directions, by digital image correlation (DIC) against an averaged image (with no strain) obtained straight after the US. DIC was performed using the MM2DPosSism subprogram of MicMac [37] with a comparison box of 5 *×* 5 pixel. An example of recovered displacements and a comparison with the input signal is shown in Fig. 2 top. The waveforms of the recovered displacements display the same carrier frequency *f*0 as the input signal, with an envelope encompassing both the main pulse, centered at *t*0*≈* 0.70 ms and an echo which, because of the stroboscopic recording, folds back at the beginning of the movie.

Analogous to the flow field estimation (see V D 1) homodyne detection was performed on the cell motion maps. This yields:

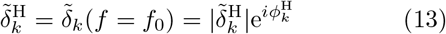

This leads to pixel-wise homodyne amplitude 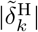 and phase 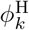 maps (see Figs. 2 and 3). These complex amplitude maps were then warped to the same shape as the fluorescence images (see V C 1) using an affine transformation with bi-linear interpolation. This led to a direct pixel-to-pixel correspondence between the images acquired with the two imaging systems.

#### 3. Homodyne strains computation

From bi-dimensional displacement maps δ*k*(*x, y*), pure and simple shear strains can be classically calculated as:

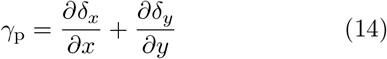

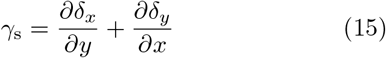

respectively.

Analogous, homodyne strains are computed from the homodyne displacements maps 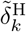 (see V D 2):

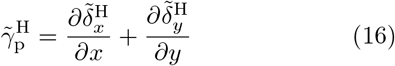

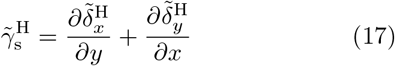

#### 4. Shear modulus derivation

By considering the components of the flow alongside *x* and *y* separately, the pure, 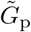, and simple, 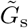, dynamic shear moduli are obtained by dividing the displacement maps (see Eq. 4) by the applied stress (see Eq. 2):

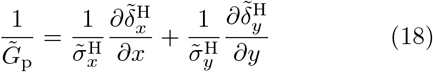

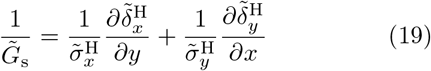

The dynamic shear moduli were computed for every area of interest and allows us to compare the rigidity of different cells and regions of cells. An example of a shear modulus map recovered from both the pure and simple strains is presented in Fig. 4. For simplicity we will use the mean of the pure and simple shears as “dynamic modulus” in the following:

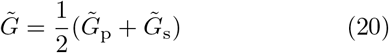

#### 5. Statistical analysis

All the distributions of dynamic shear modulus presented in this study were assessed to be non-normally distributed using the Shapiro-Wilk test [38] (pvalue *<* 0.001). Statistical differences between the tested phenotypes were hence assessed using the Mann-Whitney U test [39].

### E. Supplementary materials

Cropped videos of the cell types displayed in Fig. 2 are provided.

### F. Data and code availability

Pre-processed data alongside analysing tools needed to reproduce the results presented in this article are available from GitHub (te be uploaded).

## VI. SUPPLEMENTARY FIGURES

## Notes

### Competing Interest Statement

The authors have declared no competing interest.

